# A TSST-1 structural motif disrupts endothelial programs required for vascular regeneration

**DOI:** 10.64898/2026.05.12.724633

**Authors:** Sharon S. Tang, Oluwadamilola Babatunde, Phuong M. Tran, Xiao-Jun Wu, Annabella G. Castiglione, Kyle J. Kinney, Dhruthi Suresh, Kavi Prem Milan Mehta, Wilmara Salgado-Pabón

## Abstract

*Staphylococcus aureus* causes profound vascular damage during infection, where it inflicts vascular injury across organs and generates lesions that fail to heal. Superantigens are major virulence factors in *S. aureus* infections, yet their direct effects on vascular repair remain unclear. We provide evidence that TSST-1 disrupts endothelial regeneration through coordinated mechanisms. TSST-1 interferes with collective directed migration, impairing endothelial cell directional persistence and preventing re-endothelialization *in vitro*. These defects stem from cytoskeletal disorganization, characterized by stress fiber accumulation and loss of lamellipodia, and broad suppression of motility-associated secreted factors. In an *ex vivo* aortic ring assay, TSST-1 suppresses angiogenic sprouting and generates dysmorphic vascular networks. Proteomic profiling reveals a shift toward matrix rigidity, adhesion stabilization, and overall suppression of angiogenesis. These activities map to a conserved dodecapeptide motif. Hence, TSST-1 suppression of vascular repair may convert sites of tissue injury into persistently non-healing niches suited for *S. aureus* persistence.

## INTRODUCTION

*Staphylococcus aureus* remains a leading cause of life-threatening infections, including pneumonia, sepsis, and infective endocarditis (IE).^1–3^ *S. aureus* inflicts vascular endothelial injury and fosters lesions that fail to heal across organ systems, a hallmark of invasive staphylococcal disease.^4–9^ In IE, the bacterium rapidly assembles valve-bound vegetations composed of bacterial microcolonies interwoven with fibrin and platelets, admixed with inflammatory and necrotic debris.^10^ These friable structures fragment, seed emboli, and precipitate end-organ injury, including infarcts in solid organs driving morbidity and mortality.^11^ Beyond IE, *S. aureus* pneumonia and skin/soft-tissue infections share a signature of endothelial damage and toxin-driven necrosis that can persist despite appropriate antimicrobial therapy.^9,12,13^ Superantigens (SAgs) are central to many of these pathologies. In experimental models they are required for fulminant IE, lethal sepsis, and kidney injury, and they potentiate tissue destruction.^14,15^ SAgs are best known for provoking massive, Vβ-skewed T-cell activation and cytokine storm that can cause toxic shock syndrome and lethal shock.^16^ Critically, SAgs also act directly on vascular endothelium, dysregulating adhesion molecules, barrier integrity, and re-endothelialization.^17,18^

In fact, angiogenic repair programs that restore endothelial integrity and re-vascularize injured tissue have emerged as convergent targets of *S. aureus* virulence factors critical for initiation and expansion of vegetations. In IE, mechanical or inflammatory injury to valve endothelium creates a nidus for bacterial adhesion and rapid vegetation growth.^19^ Efficient re-endothelialization would ordinarily limit thrombus formation and microbial colonization,^20^ yet, *S. aureus* appears to actively subvert this repair process by targeting angiogenic programs via various exotoxin-driven mechanisms. Studies in the rabbit model of native-valve IE with MRSA strain MW2 demonstrate that the SAg staphylococcal enterotoxin C (SEC) is required for vegetation growth and organ injury, as SEC-deficient strains are attenuated *in vivo*.^14,21^ Strikingly, SEC retains much of its pathogenic activity when the TCR and MHC II binding sites are inactivated, revealing superantigenicity-independent functions essential for disease.^21^ Consistent with these observations, SEC has been shown to suppress microvessel formation in the *ex vivo* rabbit aortic ring model of angiogenesis and to reduce expression of endothelial VEGF-A, a master regulator of vascular repair.^21^ Additionally, SEC inhibits pentraxin-3 (PTX3), amphiregulin (AREG), and TIMP-4 production from aortic endothelial cells, converging on core motility programs and positioning anti-angiogenesis as a direct driver of failed vascular repair in IE.

Toxic shock syndrome toxin (TSST)-1, a highly prevalent SAg among mucosa-associated *S. aureus* strains, is likewise associated with invasive disease,^15,16,22^ and promotes vegetation formation in experimental IE.^15^ TSST-1 directly targets aortic endothelial cells, independently of canonical MHC II engagement, activating endothelium by upregulating adhesion molecules, increasing monolayer permeability, and impairing re-endothelialization.^17^ These effects occur in the absence of T-cell–driven inflammation, demonstrating direct modification of endothelial function. Notably, TSST-1 suppresses endothelial interleukin-8 and interleukin-6 production (chemokines essential for neutrophil recruitment), suggesting that SAgs attenuate local inflammatory responses while promoting vascular dysfunction.^17^ Collectively, the non-hematopoietic actions of SAgs provide a mechanism for sustained tissue disruption at infection sites, highlighting a direct role in vascular repair interference that is separable from their immune-activating properties.

Despite these emerging insights, the cellular mechanisms through which SAgs, such as TSST-1 and SEC, directly impair vascular repair remain poorly defined. Building on our observation that TSST-1 inhibits re-endothelialization,^17^ we sought to delineate mechanisms responsible for this impairment and consequences for angiogenic sprouting. Here we show that TSST-1 specifically disrupts directional endothelial migration by suppressing a broad network of motility-associated factors and by inducing profound reorganization of the actin cytoskeleton. Furthermore, TSST-1 inhibits microvessel sprouting and yields dysmorphic, non-productive vascular networks. Proteomic analyses identify coordinated alterations in matrix remodeling, adhesion-complex dynamics, and angiogenic signaling that converge to impair vessel formation. Guided by evidence that TSST-1 can signal non-hematopoietic cells via a conserved dodecapeptide,^23,24^ we mapped the anti-regenerative activities to this motif. Our findings place TSST-1 alongside SEC and β-toxin^25^ as part of a coordinated strategy that targets distinct but convergent nodes in the angiogenic program.

## RESULTS

### TSST-1 targets directed endothelial cell migration

Previous work showed that TSST-1 impairs re-endothelialization in an *in vitro* wound-healing assay when tested at sublethal concentrations up to 200 µg mL⁻¹ on immortalized human aortic endothelial cell (iHAEC) monolayers.^17^ Whether this defect reflected altered proliferation or migration, however, remained unresolved. To address this, we first assessed proliferative capacity using an EdU incorporation assay, which revealed no reduction in proliferation in TSST-1–treated cells (**Figure 1A**). In contrast, cell-exclusion assays demonstrated a marked decrease in gap closure in monolayers exposed to TSST-1. This migration defect persisted, and became more pronounced, when cell division was blocked with mitomycin C, indicating that impaired gap closure occurred independently of proliferation (**Figure 1B**). Together with prior findings, these data demonstrate that TSST-1 selectively disrupts endothelial cell migration without detectable effects on viability or proliferation.

**Figure 1.**
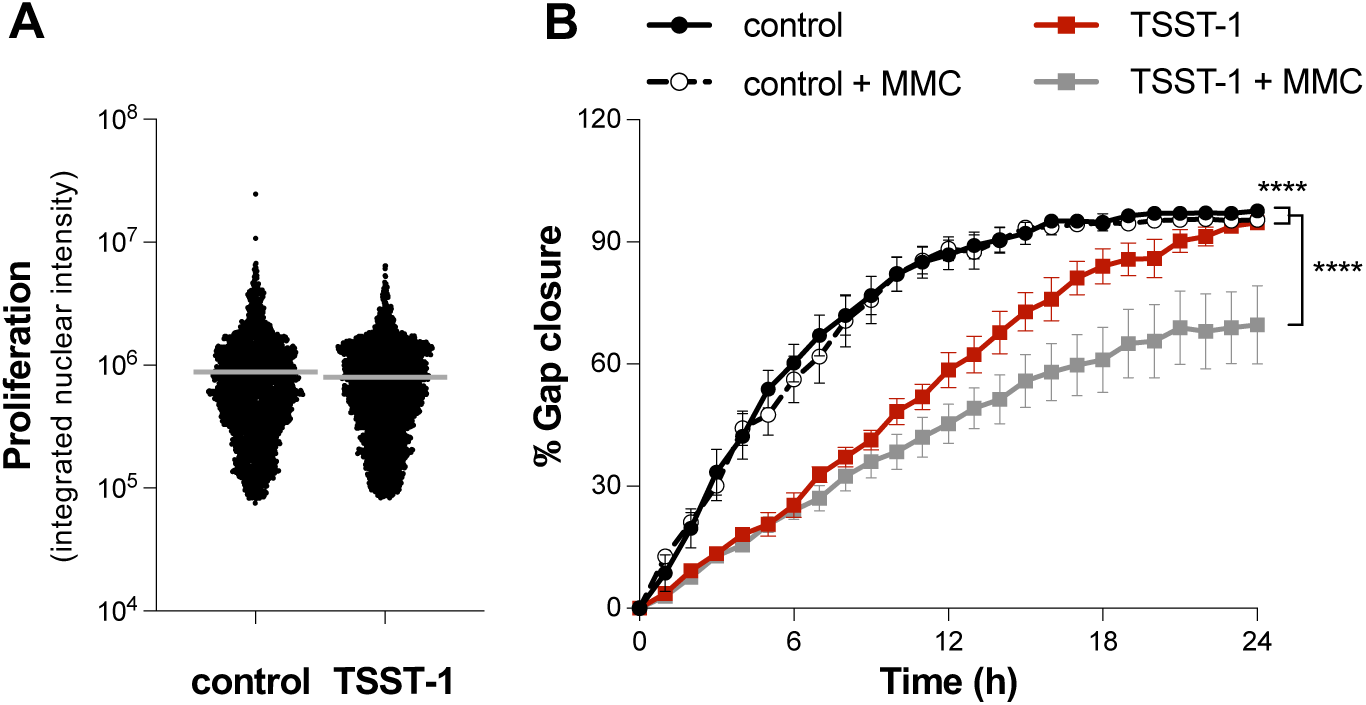
TSST-1 selectively inhibits endothelial cell migration. **(A)** Cell proliferation of iHAEC seeded at 7,000 cell/well in a 96-well plate and treated with TSST-1 (50 μg mL^⁻^¹) overnight to up to ∼80% confluency. Edu incorporation measured for 2 h in control versus TSST-1–treated cells. **(B)** Time course analysis of iHAEC grown to confluency in silicone inserts that create uniform gaps upon removal and treated with either TSST-1 (50 µg mL−1) or untreated ± mitomycin C (MMC; 2 µg mL^⁻^¹). Images acquired every 30 min for 24 h. Data represent mean ± SEM; \*\*\*\**p* < 0.0001; two-way ANOVA with Holm–Šídák’s multiple comparisons test.

To quantify how TSST-1 alters endothelial cell migration, we performed single-cell tracking using time-lapse images collected during the cell-exclusion assays. Representative ges acquired every 30 min for 24 h. Data represent mean ± SEM; \*\*\*\**p* < 0.0001; two-way ANOVA with m–Šídák’s multiple comparisons test. phase-contrast images at 0 h and 12 h show slower gap closure in TSST-1–treated monolayers compared with controls (**Figure 2A**). For quantitative analysis, individual endothelial cells were tracked across the imaging period using CellTraxx, an automated platform that identifies cell centroids, links positions across sequential frames, and reconstructs complete migration trajectories.^26^ Corresponding representative trajectory plots for control and TSST-1–treated conditions are shown in **Figure 2B**. Cell-migration behavior was evaluated using four parameters: (i) average velocity, representing instantaneous migration speed; (ii) Euclidean distance, measuring net displacement from the point of origin; (iii) forward-displacement index, measuring net advance; and (iv) directional persistence index, measuring directional movement across the entire trajectory. These are standard metrics for motility analyses and distinguish speed from directionality in an unbiased fashion.^27^

**Figure 2.**
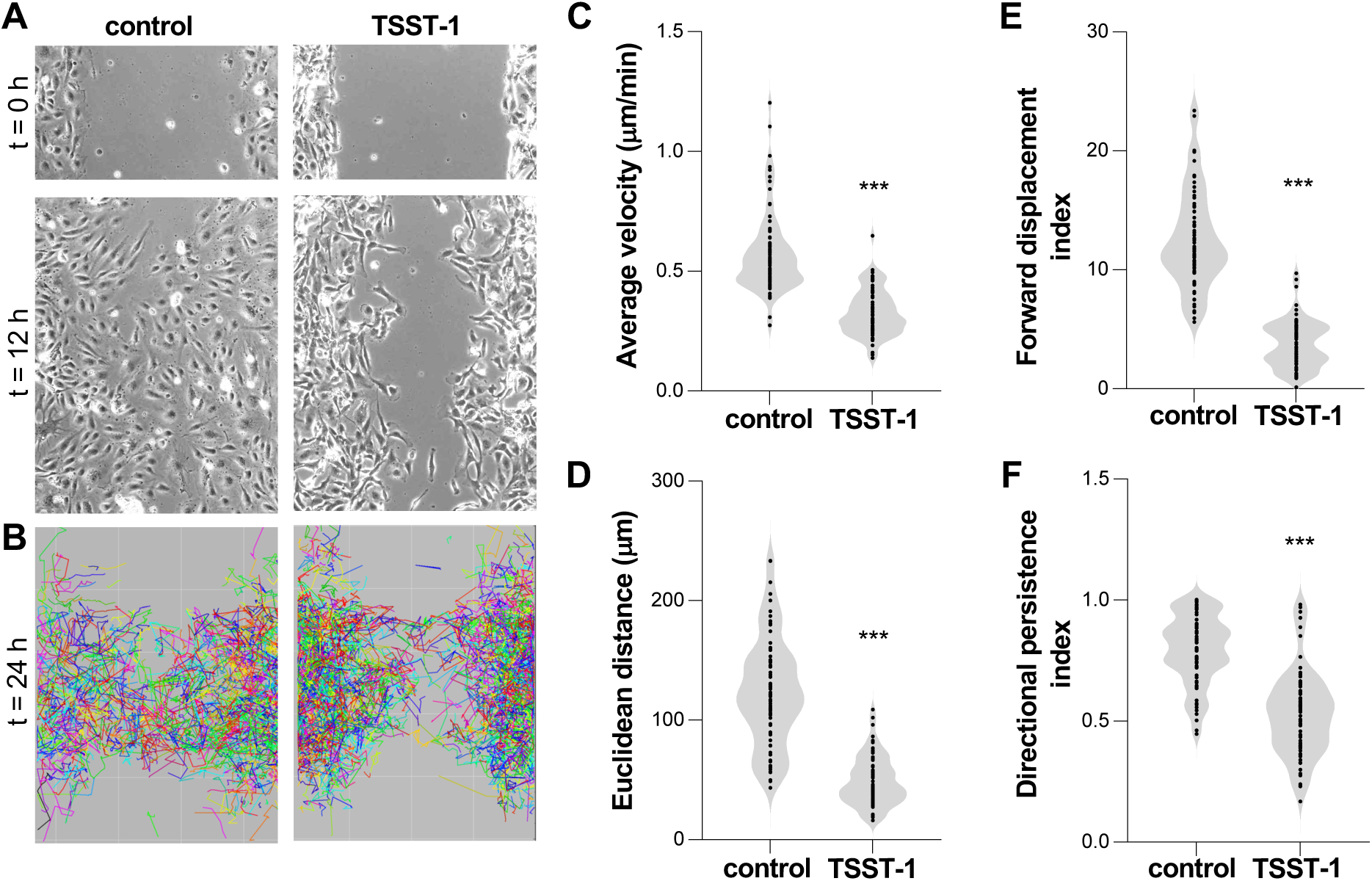
TSST-1 targets directional displacement required for effective gap closure. **(A)** Representative phase-contrast images of cell-exclusion assays at 0 h and 12 h for control and iHAEC monolayers treated with TSST-1 (50 μg mL⁻¹) . **(B)** Tracked migration paths from individual cells over 24 h. Images acquired every 30 min for 24 h. Trajectories median-filtered and drift-corrected prior to analysis. **(C)** Instantaneous velocity of a cell represented as average velocity per track (μm min⁻¹). **(D)** Euclidean distance, net displacement from the point of origin calculated as 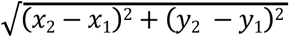. **(E)** Forward displacement index, directional efficiency calculated as (*x*_2_−*x*_1_)/total path length. **(F)** Directional persistence, directed movement toward closing the gap calculated as Euclidean distance/total distance travelled. Each symbol represents an individual cell. Data are pooled from 3 independent experiments, shown as violin plots; \*\*\**p* < 0.001. Non-parametric one-way ANOVA, Mann-Whitney test.

Control monolayers displayed rapid and directionally consistent migration. Average velocities were high (0.58 ± 0.02 μm min⁻¹; mean ± SEM) and cells achieved substantial Euclidean displacement (124 ± 5.2 μm) with correspondingly elevated forward-displacement index (12.6 ± 0.5) and directional persistence index (0.8 ± 0.02), indicating effective traversal across the exclusion gap (**Figures 2C–2F**). In TSST-1–treated monolayers, migration parameters were collectively reduced (velocity, 0.33 ± 0.01 μm min⁻¹; Euclidean displacement, 49 ± 2.4 μm; forward-displacement index, 3.7 ± 0.2; directional persistence index, 0.5 ± 0.02). Collectively, these quantitative measurements show that TSST-1 treatment is associated with a migratory state in which cells attempt to move but fail to generate the coordinated, directional displacement required for effective gap closure.

### TSST-1 disrupts consolidation and stabilization of the leading edge in migrating endothelial cells

TSST-1 alters cell migration patterns in a manner that suggests disruption of cytoskeletal or polarization mechanisms required for persistent, directed movement. To characterize these altered migration patterns, we examined individual and collective cell morphology in time-lapse images. Under basal conditions, iHAEC monolayers advanced with a continuous and cohesive leading front marked by iHAEC with broad lamellipodia, close lateral alignment, and uniform forward orientation into the gap (**Figure 3A**). In TSST-1–treated monolayers, the leading edge appeared irregular, with fragmented and discontinuous margins (**Figure 3B**). Individual cells frequently exhibited multipolarity and reduced formation of broad lamellipodial sheets, instead displaying short extensions and highly branched protrusions in multiple directions. Trailing-edge morphology was also altered, with elongated rear regions and persistent tethers commonly observed. These single-cell features are consistent with reduced front continuity, diminished coordination among neighboring cells, and limited net advancement over time compared with control monolayers.

**Figure 3.**
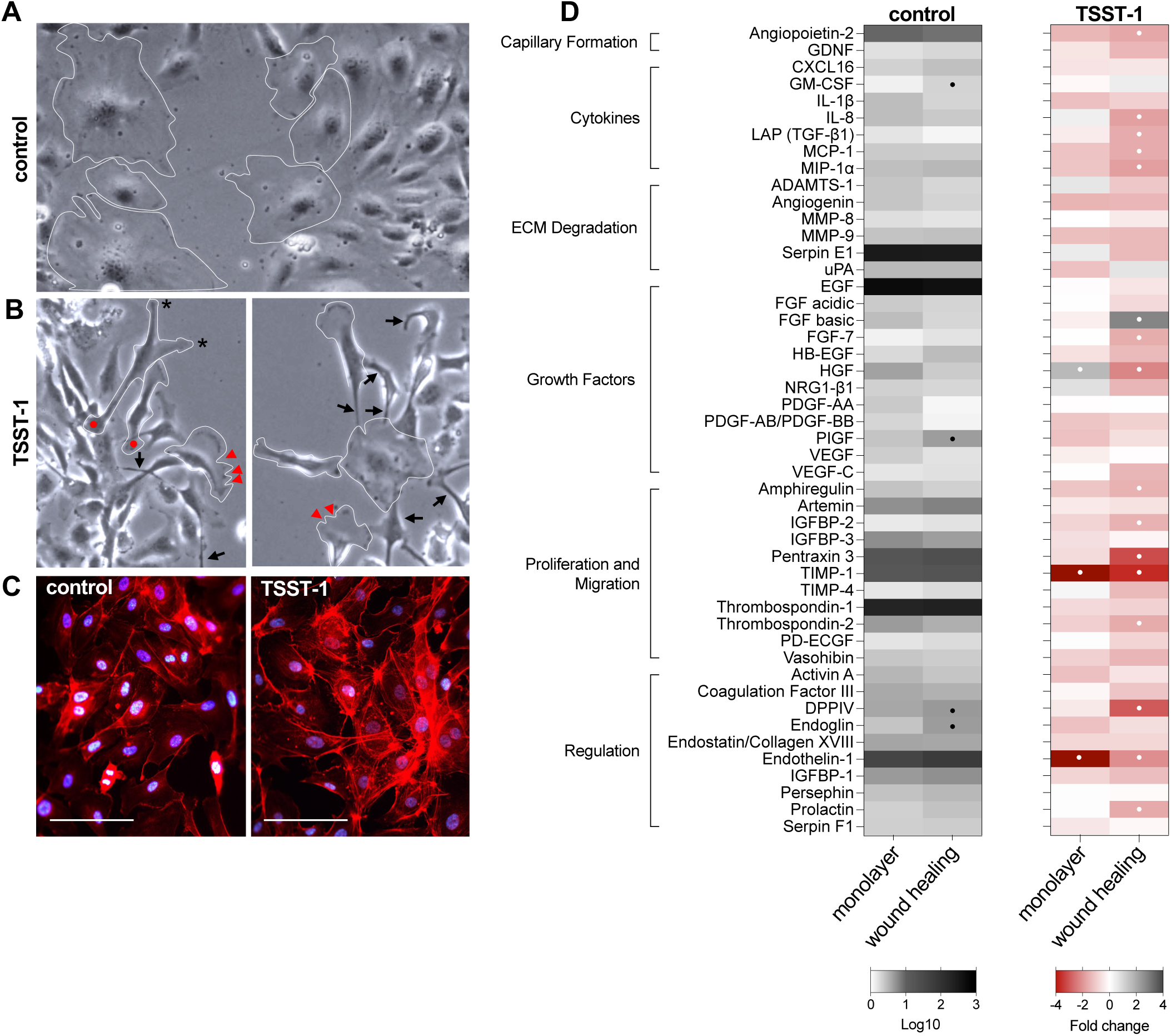
TSST-1 disrupts leading-edge stabilization and suppresses pro-migratory secretome, polarity, and adhesion programs. **(A) Cell-exclusion assay, untreated.** Representative phase-contrast images of leading-edge monolayers migrating into the gap. Cell outlines highlight iHAEC with broad lamellipodia, close lateral coordination, and uniform forward orientation toward the wound. **(B) Cell-exclusion assay, TSST-1 (50 μg mL**^⁻^**¹).** Representative phase-contrast images showing disrupted leading edges characterized by fragmented, discontinuous margins. Cell outlines reveal iHAEC with reduced lamellipodial sheets and abnormal protrusive behavior, including multipolar morphology (asterisk), short extensions (red arrowhead), highly branched, tortuous protrusions projecting in multiple directions (arrows), and elongated rear regions with persistent tethers (red dot). **(C)** iHAEC seeded at semi-confluency on fibronectin-coated glass ± TSST-1 (6 μg mL^⁻^¹) for 4 h. Actin cytoskeleton visualized with Acti-Stain 555 phalloidin (red); nuclei counterstained with DAPI (blue). Scale bar, 100 μm. **(D) Proteome Profiler Human Angiogenesis Array.** *Monolayer:* iHAEC grown to near-confluency on gelatin-coated plates ± TSST-1 (50 μg mL^⁻^¹) for 24 h. *Wound healing (cell-exclusion assay):* iHAEC cultured to confluency in silicone inserts to generate uniform gaps ± TSST-1 (50 μg mL^⁻^¹) for 24 h. Control data represent mean absolute values. Treated data represent fold change relative to controls from three independent experiments. (•) Angiogenesis-related factors showing >1.5-fold increase or decrease. In controls, wound-healing versus monolayer. In treated conditions, TSST-1–treated monolayer to monolayer control, and TSST-1–treated wound healing to wound healing control.

To evaluate the effects of TSST-1 on cytoskeletal organization, iHAECs were seeded at semi-confluent density on fibronectin-coated glass slides and examined by F-actin immunofluorescence. Untreated monolayers displayed uniform cortical actin and well-defined intercellular junctions. In contrast, TSST-1 exposure produced a marked reorganization of the actin cytoskeleton (**Figure 3C**). Cells exhibited prominent and elongated stress fibers, extensive F-actin bundling, and numerous thin protrusive actin extensions emerging from multiple regions along the cell perimeter.^28^ Hence, TSST-1 impairs directed collective migration through alterations in lamellipodia architecture and actin dynamics.

### TSST-1 suppresses the pro-migratory secretome and disrupts polarity-adhesion programs

To examine soluble factors associated with the morphological and cytoskeletal changes observed in TSST-1–treated iHAEC, we used a human angiogenesis proteome array to profile 48 secreted proteins in either confluent monolayers or actively migrating cells in the presence or absence of TSST-1 (50 µg mL⁻¹) (**Table S1**). This approach distinguished signals present under basal junction-maintaining conditions from those associated with active protrusion and collective migration. Relative to confluent monolayers, migrating cells showed increased levels of GM-CSF, PlGF, DPPIV, and Endoglin (**Figure 3D**), demonstrating their activated, reparative, and pro-angiogenic state. In TSST-1–treated monolayers, levels of TIMP-1 and Endothelin-1 (ET-1) were reduced relative to untreated controls (**Figure 3D**). This reduction is consistent with dysregulated pericellular proteolysis and impaired focal-adhesion turnover.^29^ These decreases also help explain the cytoskeletal alterations observed by F-actin immunofluorescence.

TSST-1–treated migrating iHAECs exhibited a broadly reduced secretory profile across multiple categories of motility-associated factors. Proteins linked to lamellipodial activity (IL-8, MCP-1, MIP-1α, HGF, AREG, FGF-7), polarity and junctional regulation (Ang-2, TGF-β1, IGFBP-2, PTX3, PRL), adhesion and extracellular-matrix remodeling (TIMP-1, TSP-2, DPPIV), and regulators of contractile tone (ET-1) were all decreased relative to untreated migrating cells (**Figure 3D**).^29^ In contrast to these widespread reductions, FGF basic (FGF2) was increased in TSST-1–treated cells. As a classic stress-induced survival factor, elevated FGF2 likely reflects a compensatory response to the loss of trophic cues and may represent a nonspecific attempt to sustain motility under conditions of impaired cytoskeletal organization and diminished directional guidance.^30^ Together, these secretome alterations point to a coordinated collapse of the endothelial migration program.

### TSST-1 induces a suppressive and dysmorphic sprouting phenotype in rabbit aortic ring explants

Angiogenesis remodels the vascular network through the growth of new capillaries from pre-existing vessels. Pathogens that modulate angiogenesis typically do so by targeting multiple steps of this process.^31^ To evaluate the impact of TSST-1 in a physiologically relevant context, we employed the *ex vivo* rabbit aortic ring assay, a gold-standard model that preserves the native vessel microenvironment. Thoracic and abdominal aortic segments (∼1 mm width) from New Zealand White rabbits were embedded in a thin layer of GFR-Matrigel to stimulate microvessel sprouting from the cut edge. In untreated controls, both ring types generated robust radial sprouting that increased in density and complexity over 14 days. In contrast, TSST-1 inhibited sprouting in a dose-dependent manner (**Figure 4A**). Low concentrations (<10 µg mL⁻¹) had minimal effects, with sprouting areas comparable to controls. Treatment with 10–20 µg mL⁻¹ TSST-1 reduced microvascular outgrowth by approximately 25%, whereas 50 µg mL⁻¹ nearly abolished angiogenic activity. Representative images of rings exposed to the submaximal 20 µg mL⁻¹ dose demonstrate the marked loss of microvessel formation and highlighted the distinct sprouting morphology induced by TSST-1 (**Figure 4B**).

**Figure 4.**
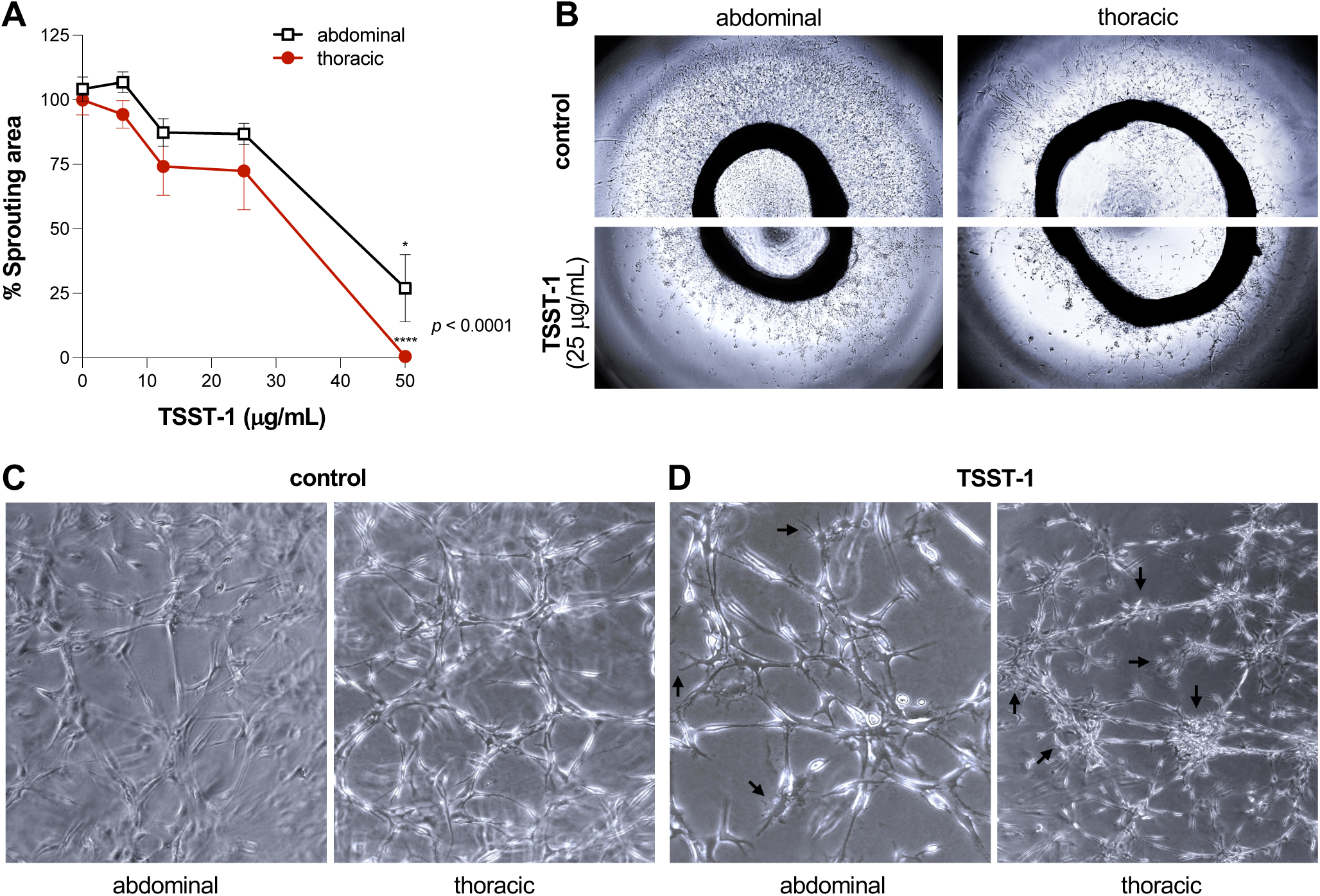
TSST-1 reduces angiogenic branching and disrupts structural organization in the rabbit aortic ring model of angiogenesis. Thoracic and abdominal aortic ring explants from New Zealand White rabbits cultured on GFR-Matrigel for 14 days in the presence or absence of TSST-1. **(A)** Quantification of sprouting area in aortic rings treated with increasing concentrations of TSST-1. Data represent mean ± SEM. \*\*\*\**p* < 0.0001 by two-way ANOVA with Holm–Šídák’s multiple comparisons test. *p-*values indicate comparisons between treated and untreated conditions. **(B)** Phase-contrast microscopy of abdominal and thoracic aortic rings imaged at day 14. Microvessels extending from untreated rings (top) form continuous sprouts, while rings treated with TSST-1 (25 μg mL^⁻^¹; bottom) show visibly reduced and irregular outgrowth. **(C–D)** Phase-contrast images illustrating the effect of TSST-1 (25 μg mL^⁻^¹) on vascular network morphology at day 14. **(C)** Untreated rings exhibit typical healthy angiogenic sprouting characterized by a dense and highly interconnected network of elongated structures. Branch points are frequent and relatively evenly distributed, and sprouts extend in smooth, continuous trajectories with aligned cells forming coherent cords. **(D)** Rings treated with TSST-1 show dysmorphic networks with reduced cohesion. Sprouts are shorter and thicker, and branching is more irregular, forming localized clusters and star-like arrangements instead of continuous cords. Abdominal: arrows highlight terminal sprout ends with extended thin, tapered protrusions consistent with numerous tip cell filipodia. Thoracic: arrows point to dense radial clusters or abrupt, star-like formations suggestive of impaired directional extension.

To further characterize the impact of TSST-1 on angiogenic morphogenesis, we examined the architecture of the sprouting networks generated by abdominal and thoracic aortic rings. Without treatment, aortic rings exhibited the expected angiogenic morphology, consisting of dense multicellular sprouts that extended radially from the explant and formed a highly interconnected microvascular plexus. Sprouts appeared continuous, aligned, and organized into cohesive cord-like structures typical of robust endothelial outgrowth (**Figure 4C**). Submaximal TSST-1 exposure generated aberrant sprouting phenotypes with markedly reduced branch density. Sprouts frequently appeared fragmented, more tortuous, and forming dense clusters or knots rather than smooth radial extensions. Many sprouts displayed a jagged, spiky appearance forming filopodia-like extensions extending from central cellular masses consistent with disrupted tip cell morphology and polarity (**Figure 4D**).^32^ Thus, depending on exposure levels, TSST-1 suppresses angiogenic sprouting or causes the formation of dysmorphic vascular networks.

### TSST-1 induces proteomic shifts that promote matrix rigidity, adhesion stabilization, and stress responses

To define the TSST-1–induced molecular pathways underlying its anti-angiogenic activity, we performed LC-MS/MS–based proteomic profiling on aortic explants cultured for 14 days in the presence or absence of TSST-1 (50 μg mL^−1^). After applying a threshold of ≥10 total spectral counts across all samples and removing proteins detected in matched culture media (**Table S2**), we identified 182 proteins in sprouted aortas and 166 proteins in TSST-1–treated aortas. Of these, 147 proteins were shared between conditions, whereas 35 were unique to sprouted aortas and 19 to TSST-1–treated aortas. These curated protein datasets were analyzed using the Metascape platform to identify pathways and biological processes affected by TSST-1 (**Table S3**).^33^ Pathway enrichment analysis and quantification of protein abundance across functional groups demonstrated that TSST-1 exposure diminished programs associated with cytoskeletal regulation, contractility, and motility (cluster 2), and vessel wall activation during angiogenesis (cluster 5) (**Figures 5A and 5B**). Moreover, the proteome of TSST-1–treated aortas were diminished in pathways associated with the ITGA1-ITGB1-COL6A3 complex and positive regulation of lamellopodium organization, while enriched for pathways associated with protein homeostasis (cluster 4) and for proteins in the localization of PINCH-ILK-PARVIN to focal adhesion pathway (**Figure 5A**).

**Figure 5.**
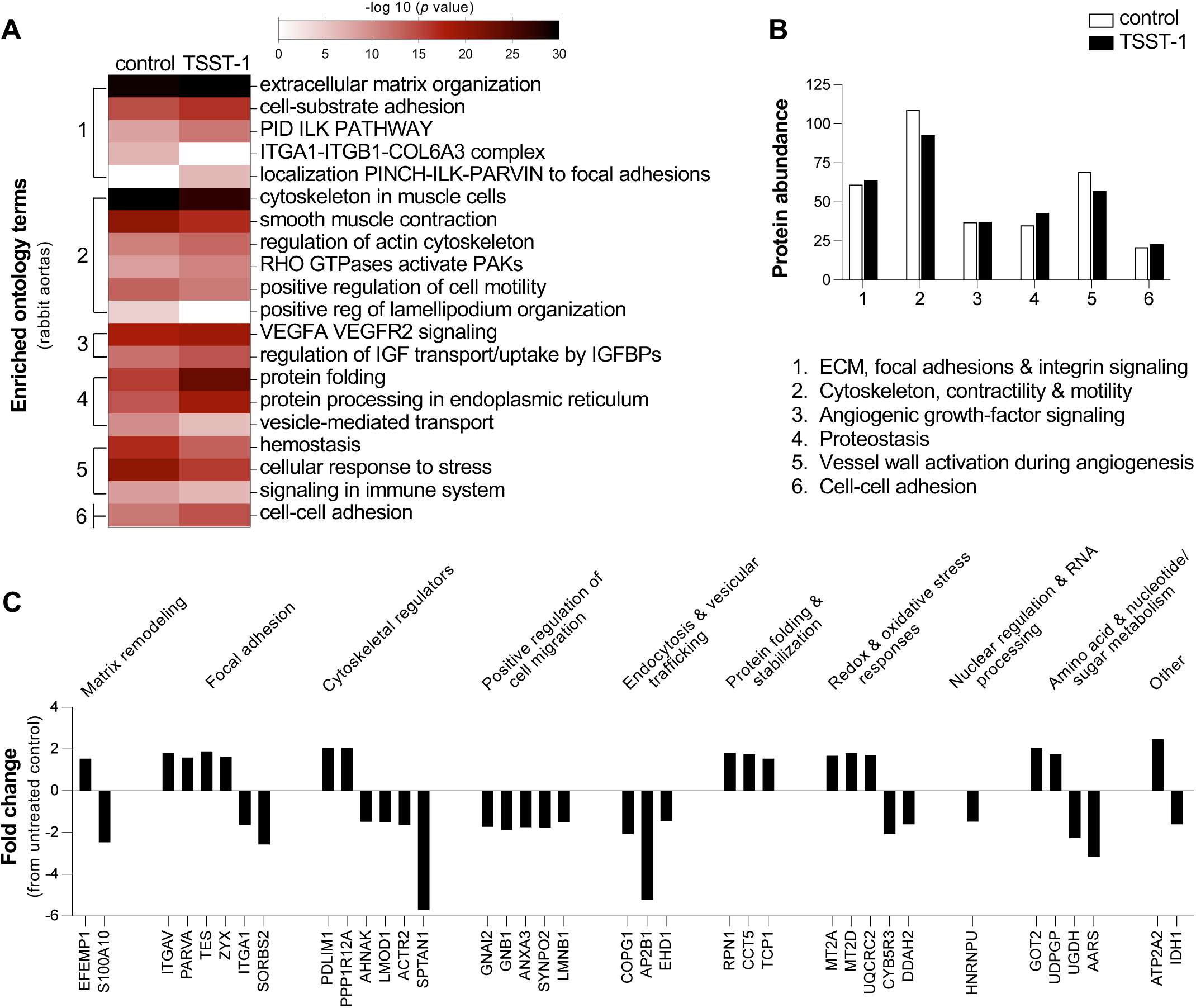
TSST-1 alters angiogenic, cytoskeletal, ECM, and stress-response pathways in aortic ring explants. Abdominal and thoracic aortic explants cultured for 14 days ± TSST-1 (50 μg mL^⁻^¹), followed by LC-MS/MS proteomic profiling of aortic rings. Changes delineate the TSST-1–responsive aortic proteome associated with inhibition of angiogenic sprouting. **(A) Pathway enrichment analysis of aortic ring proteomes.** Curated datasets analyzed using Metascape. Heatmap shows significantly enriched Gene Ontology terms clustered into biological categories. Color intensity reflects −log10 (*p*-value). Identified proteins were grouped into six functional clusters. **(B) Protein abundance by functional category.** Protein abundance across functional clusters in aortic rings for control versus TSST-1–treated aortas. Highlights detection of specific proteins across conditions to identify pathways that were depleted or enriched. **(C) Differential enrichment of proteins in response to TSST-1.** Proteins from TSST-1–treated explants with >1.5-fold change in total spectral counts compared to controls. Proteins organized by functional group.

To identify protein changes specifically associated with TSST-1–mediated inhibition of angiogenic sprouting, we compared proteins exhibiting a >1.5-fold change (increase or decrease) in total spectral counts between TSST-1–treated aortic rings and sprouted controls (**Figure 5C**). We found that TSST-1 increased abundance of ECM-structuring protein EFEMP1 and decreased abundance of S100A10, a component of the ANXA2/S100A10 complex that localizes tissue-type plasminogen activator and plasminogen at the cell surface. These changes could alter the pericellular proteolysis landscape.^34–36^ TSST-1 also modified focal-adhesion composition, inducing a shift in integrin representation from ITGA1 to ITGAV. Additionally, proteins associated with adhesion-complex components (PARVA, TES, and ZYX) were elevated, whereas the adaptor SORBS2, a negative regulator of excessive adhesion signaling, was reduced.^36–38^ Overall, this adhesion and proteolysis signature indicates that TSST-1 promotes over-stabilized, poorly tunable adhesions.

These changes were accompanied by coordinated alterations in cytoskeletal regulators. The contractility-associated regulatory subunit PPP1R12A was increased, consistent with enhanced modulation of actomyosin activity.^101^ In contrast, components that support branched actin assembly and cortical integrity, including ACTR2 and SPTAN1, were reduced, suggesting a weakened lamellipodial and membrane–cytoskeleton interface.^39–41^ Additional reductions of LMOD1, an actin nucleator that promotes filament elongation, and AHNAK, a scaffold essential for cortical organization and membrane coupling, further indicate disruption of filament architecture and peripheral stability.^34,35,100^ Conversely, the actin–adhesion scaffold PDLIM1 was increased, consistent with a shift toward stress-fiber–enriched structural programs and reinforced adhesion complexes. In parallel, multiple factors that support cell migration (GNAI2, GNB1, ANXA3, SYNPO2, LMNB1) were reduced in TSST-1–treated aortas, as were components involved in endocytosis and membrane recycling (COPG1, AP2B1, EHD1) relative to controls.^42–45^ Collectively, this proteomic signature indicates coordinated changes across cytoskeletal organization, adhesion/migration machinery, and membrane trafficking in TSST-1–treated aortic rings that are consistent with impaired directed migration and dysmorphic angiogenic sprouting.

### TSST-1 causes a secretome-wide shift toward anti-angiogenic signaling and extracellular matrix stabilization

To fully capture the aortic response to TSST-1, we profiled cell-free conditioned media from aortic rings cultured in the presence or absence of TSST-1. The same analytical workflow and filtering thresholds applied to the aortic ring datasets were used for these samples. We identified 53 proteins in sprouted aortas and 52 proteins in TSST-1–treated aortas. Of these, 46 proteins were shared between conditions, whereas 7 were unique to sprouted aortas and 6 to TSST-1–treated aortic cultures (**Table S3**). Pathway enrichment analysis and quantification of protein abundance across functional groups demonstrated that TSST-1 treatment enriched for proteins in pathways associated with supramolecular fiber organization (cluster 4), and cell adhesion and junction organization (cluster 6), while depleted those in platelet activation, complement and coagulation pathways (cluster 5) (**Figures 6A and 6B**).

**Figure 6.**
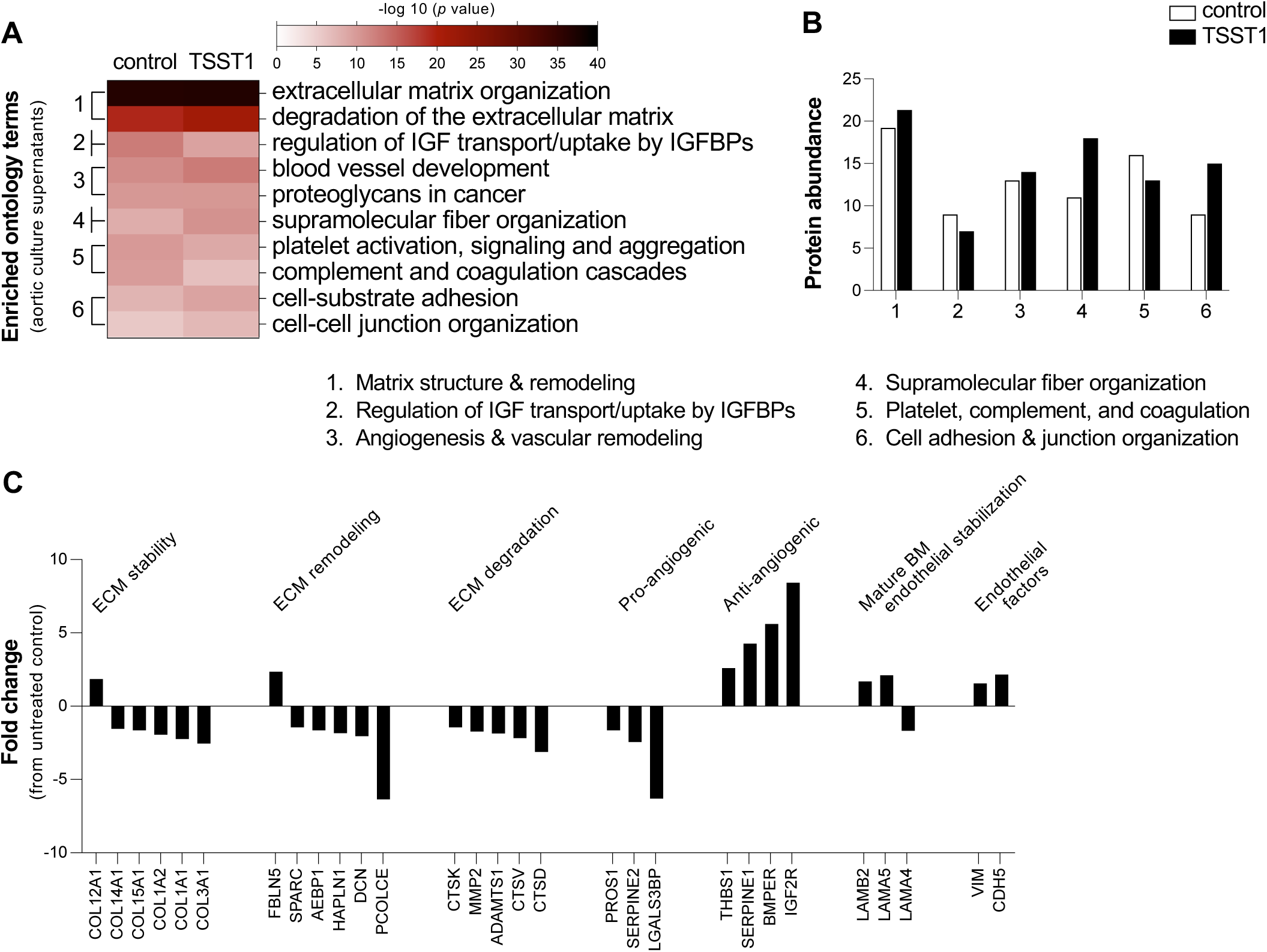
TSST-1 remodels the secreted proteome of aortic ring explants. Abdominal and thoracic aortic explants cultured for 14 days ± TSST-1 (50 μg mL⁻¹), followed by LC-MS/MS proteomic profiling of conditioned media. Changes delineate the TSST-1–responsive secreted proteome associated with inhibition of angiogenic sprouting. **(A) Enriched pathways in conditioned media proteomes.** Curated datasets analyzed by Metascape. Heatmap displays significantly enriched Gene Ontology terms clustered into major biological categories. Color intensity reflects −log 10(p-value). Identified proteins were grouped into 6 functional clusters. **(B) Functional classification of soluble proteins.** Protein abundance across functional clusters in conditioned media for control versus TSST-1–treated aortas. Highlights detection of specific proteins across conditions to identify pathways that were depleted or enriched. **(C) Differential enrichment of proteins in response to TSST-1.** Proteins from conditioned media of TSST-1–treated explants with >1.5-fold change in total spectral counts compared to controls. Proteins organized by functional categories.

Analysis of proteins showing >1.5-fold changes from untreated controls revealed that TSST-1 induced a pronounced anti-angiogenic secretome signature (**Figure 6C**). Proteins that sequester growth factors such as VEGF, BMP, and IGF, limit pericellular proteolysis, and suppress endothelial motility were markedly increased (THBS1, SERPINE1, A2M, IGF2R, and BMPER). ^46–49^ In parallel, several pro-angiogenic supports were depleted (PROS1, SERPINE2, and LGALS3BP).^50–53^ This signature is consistent with an extracellular environment prone to growth-factor sequestration, reduced proteolysis, and impaired endothelial invasion.

Matrix-associated pathways also showed broad shifts toward reduced remodeling activity. Multiple degradative enzymes, including MMP2, ADAMTS1, and several cathepsins (CTS), were decreased in TSST-1–treated aortic rings.^34,54–60^ Non-degradative remodeling components changed in abundance, with increases in elastic fiber- and fibril-associated proteins such as FBLN5 and COL12A1, accompanied by decreases in small leucine-rich proteoglycan DCN, hyaluronan-linking protein HAPLN1 (components that produced a hydrated, highly regulated matrix), and fibrillar collagens I and III and associated linker proteins.^61–65^ This signature is indicative of a move toward a stiffer or more organized extracellular matrix. Basement membrane (BM)–associated proteins also shifted, with LAMA4 decreased and LAMA5 and LAMB2 increased, indicating a BM composition typical of mature BM that stabilizes the endothelium.^66–68^

Finally, the endothelial-related proteins soluble VE-cadherin (CDH5) and vimentin (VIM) were elevated in the medium. These results are consistent with junctional breakdown and cytoskeletal stress. This signature aligns with our previous observation that TSST-1 decreases total VE-cadherin and disrupts its junctional distribution, indicating that TSST-1 compromises both adherens-junction integrity and endothelial homeostasis.^17,69,70^ Altogether, the extracellular protein changes demonstrate TSST-1–associated alterations across ECM-remodeling enzymes, fibrillar and BM components, and endothelial-derived factors.

### The TSST-1 dodecapeptide mediates novel anti-migratory and anti-angiogenic activities

SAgs interact with multiple cell types through a conserved dodecapeptide located at the base of the central α-helix (**Figure 7A**).^16^ This region has been shown to mediate epithelial engagement and translocation across mucosal barriers.^23,24,71^ To determine whether TSST-1–mediated inhibition of cell migration and angiogenesis depends on this conserved motif, we generated a panel of alanine-substitution variants across the dodecapeptide (F119A, D120A, K121A, K122A, Q123A, L124A, A125S, I126A, S127A, T128A, L129A, and D130A), replacing the native alanine at position 125 with serine. Variant D130A was excluded from subsequent analyses due to insufficient protein yield for spectral measurements. Circular dichroism (CD) spectroscopy was performed on wild-type TSST-1 and the 11 analyzable variants to assess global structural integrity. Across all mutants, the magnitude and overall shape of the CD spectra closely matched wild-type TSST-1. The averaged spectrum overlapped with each individual mutant trace, and spectral variance remained within experimental noise. None of the substitutions produced significant shifts in band minima, loss of ellipticity at 208 or 222 nm, or increases in random-coil signal (**Figure 7B**). These data indicate that the substitutions tested did not alter the global secondary structure of TSST-1.

**Figure 7.**
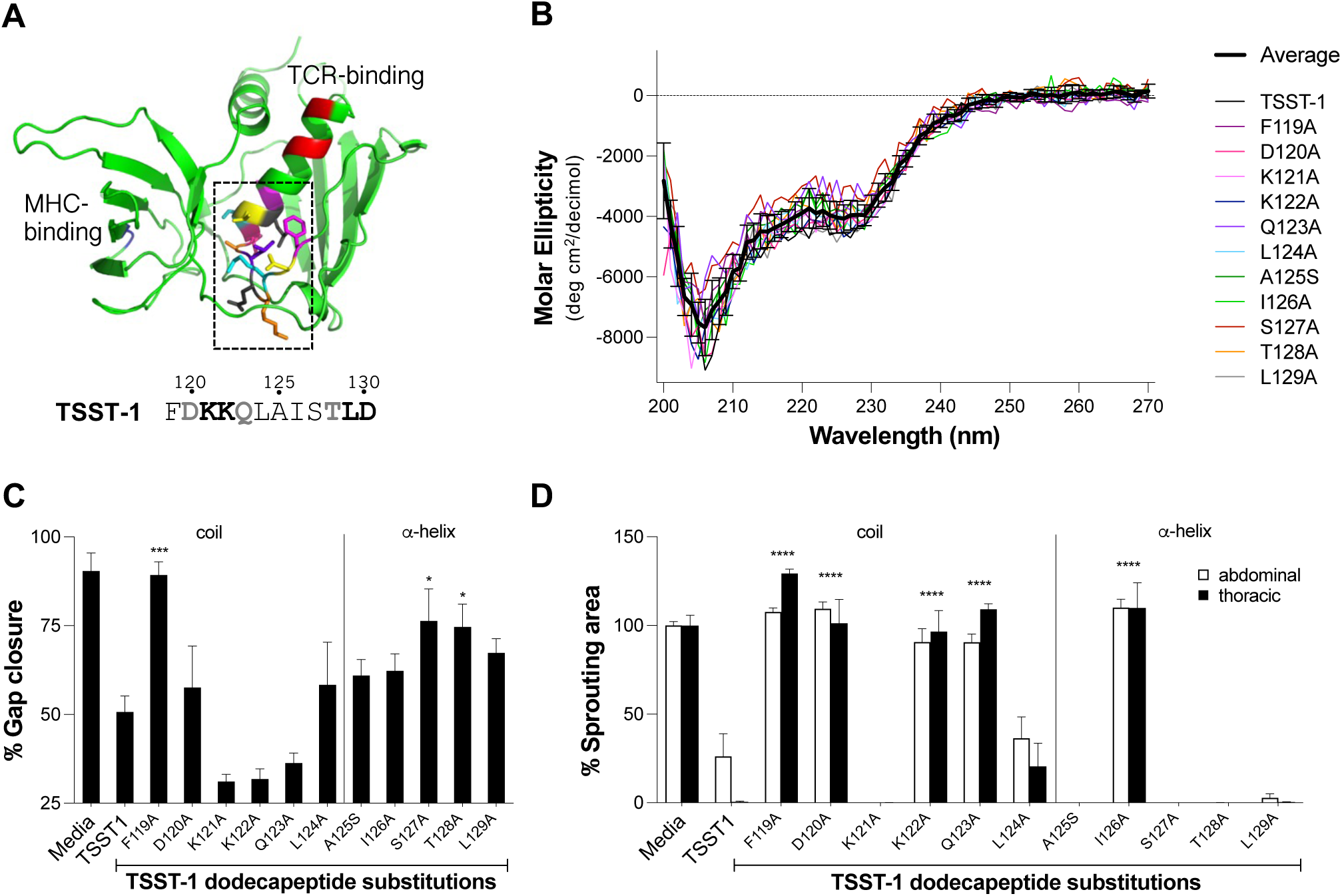
The TSST-1 dodecapeptide mediates the suppressive vascular regeneration activities. Specific residues within the TSST-1 dodecapeptide essential for its anti-migratory and anti-angiogenic functions. **(A) Location of the TSST-1 dodecapeptide.** TSST-1 structure highlighting the conserved dodecapeptide (residues 119–130) positioned at the base of the central α-helix (colored side chains within dashed box). **(B) Secondary structure analysis of dodecapeptide variants.** Circular dichroism (CD) spectra for wild-type TSST-1 and 11 analyzable alanine-substitution variants (native alanine 125 replaced with serine). Individuals and average CD spectra shown. **(C) Functional contribution of dodecapeptide residues to inhibition of cell migration.** Time course analysis of iHAEC grown to confluency in silicone inserts that create uniform gaps upon removal ± TSST-1 or individual alanine-substitution variants (20 µg mL^⁻^¹). Images acquired every 30 min for 24 h. Data represent 20 h time point, mean ± SEM; \**p* < 0.02, \*\*\**p* = 0.0003 compared to TSST-1; Fisher’s LSD test. **(D) Contribution of dodecapeptide residues to inhibition of angiogenic sprouting.** Thoracic and abdominal aortic ring explants from New Zealand White rabbits cultured on GFR-Matrigel for 14 days ± TSST-1 or individual alanine-substitution variants (50 µg mL^⁻^¹). Quantification of sprouting area. Data represent mean ± SEM. \*\*\*\**p* < 0.0001 compared to TSST-1; two-way ANOVA with Holm–Šídák’s multiple comparisons test.

To assess whether individual residues within the TSST-1 dodecapeptide contribute to suppression of cell migration, we performed cell-exclusion assays in the presence of wild-type TSST-1 or dodecapeptide variants over a 20 h period (**Figure 7C**). The F119A variant, located within the coil region of the dodecapeptide, did not reduce migration and showed gap-closure levels comparable to untreated conditions. Variants containing substitutions within the α-helical portion of the dodecapeptide, S127A and T128A, also significantly restored gap closure, although both maintained a 14 –15% reduction compared with untreated controls. These data identified F119 as critical for TSST-1’s anti-migratory activity.

To determine whether residues within the TSST-1 dodecapeptide contribute to inhibition of angiogenic sprouting, we tested the TSST-1 variants in the aortic ring assay (**Figure 7D**). Several substitutions within the coil region of the dodecapeptide (F119A, D120A, K122A, and Q123A) restored sprouting to levels comparable to untreated controls. Among the α-helical substitutions tested, only the I126A variant restored sprouting. These results show that individual residues within the dodecapeptide differentially affect cell migration and angiogenesis, with F119 emerging as the sole residue contributing to both. Still, TSST-1 relies on the same structural motif to inhibit vascular regeneration, underscoring a robust control of movement-related processes.

## DISCUSSION

*S. aureus* deploys a broad repertoire of virulence factors that injure the vasculature, promoting tissue necrosis and impairing reparative angiogenesis.^6,9^ Although staphylococcal SAgs are classically defined by immune activation, accumulating evidence indicates that they also act directly on non-hematopoietic tissues through novel mechanisms of vascular disruption. Here, we delineate extracellular and cell-intrinsic pathways from cells and tissues with TSST-1–induced impairments in vascular regeneration. We found that TSST-1 attenuates cellular motility and disrupts polarity control. TSST-1 perturbs pathways governing adhesion turnover and matrix remodeling, thereby generating a microenvironment that is unfavorable for regenerative processes. Structure–function mapping identified a dodecapeptide coil/loop motif as the principal determinant of these effects, providing a tractable interface to disrupt SAg–endothelium interactions.

The vascular endothelium maintains barrier integrity through continual renewal, a process requiring coordinated collective migration to reseal sites of injury.^72,73^ TSST-1 impairs this function by rewiring the migratory apparatus. Single-cell tracking demonstrates a marked loss of directional persistence, accompanied by disrupted front coherence, fragmented leading edges, multipolar protrusions, elongated tethers, and prominent stress-fiber accumulation.^74^ The shift from cortical actin to thick cytoplasmic bundles mirrors endothelial responses to pathological mechanical or inflammatory cues, where excess contractile tension suppresses lamellipodial protrusion and distorts actomyosin–Arp2/3 balance.^75–78^ This combination, reinforced contractile fibers and diminished broad lamellipodia, explains the observed ineffective migration.^79,80^ Collectively, the cells generate force but cannot translate it into displacement, as contractile elements override protrusive structures and polarity cues necessary for persistent endothelial migration.^81–83^

The proteomic signature of TSST-1–treated migrating cells provide molecular context for their impaired behavior. Broad reductions in lamellipodial drivers (IL-8, HGF, AREG), polarity regulators (Ang2, IGFBP2), and adhesion-remodeling factors (TIMP1, THBS2, DPPIV) indicate a coordinated collapse of the migratory program.^29^ Notably, loss of TIMP1, which promotes FAK/Src/PI3K-dependent motility, removes a key axis of focal-adhesion turnover. ET-1 (a regulator of vascular tone that promotes directional migration and adhesion turnover) was decreased and explains the disorganized actomyosin bundles observed on fibronectin. Such defects are predicted to hinder trailing-edge release which was observed in TSST-1-treated cells.^84^ Together, these changes show that TSST-1 actively reprograms the extracellular milieu into an anti-migratory state, reinforcing cytoskeletal disorganization and adhesion malfunction.

The *ex vivo* aortic ring assay preserves a three-dimensional, multicellular vascular niche, enabling quantitative analysis of sprouting in a near-physiologic context.^85^ Beyond suppression, TSST-1 redirected morphogenesis toward fragmented, tortuous, multipolar sprouts rather than the cohesive cords observed in controls, a pattern consistent with dysregulated matrix–mechanical cues and disorganized neovessel formation.^86,87^ Proteomic profiling provides a mechanistic basis for this phenotype. Tissue-level changes reflected over-stabilized adhesions and reduced pericellular proteolysis, alterations known to impede endothelial invasion and bias sprouts toward pathological architectures through mechano-transcriptional pathways such as VEGFR2/Notch crosstalk and defective matrix remodeling.^88–90^ Concurrently, the matrix shifted toward a basement-membrane–like, quiescence-reinforcing state, and the secretome revealed a coordinated anti-angiogenic program.^91^ Together, these extracellular and cell-intrinsic alterations render the sprouting environment structurally incompatible with productive angiogenesis, explaining the dysmorphic, multipolar sprouts characteristic of TSST-1 treatment.

At the cell–matrix interface, the integrin repertoire shifted from ITGA1 (collagen/laminin binding) toward ITGAV-containing heterodimers (fibronectin/vitronectin binding). This shift indicates greater reliance on RGD-binding integrins (e.g., αvβ3/αvβ5), consistent with enhanced adhesion maturation and force transmission, and coincided with increases in zyxin and enrichment of PARVA (ILK–PINCH–PARVIN) and TES.^36–38^ Zyxin is a mechanosensitive focal-adhesion/stress-fiber scaffold that reinforces actin bundles and stabilizes adhesions under load. Increases favor mature but poorly tunable adhesions with slower turnover, conditions that limit forward endothelial movement and orderly sprout extension. In parallel, decreases in SORBS2, a scaffold/adaptor implicated in restraining excessive adhesion signaling, aligns with this over-stabilization model.^92,93^ Furthermore, increased EFEMP1 with concomitant decreases in S100A10, a cell-surface plasminogen receptor required for efficient pericellular plasmin generation, and reduced expression of matrix-degrading enzymes (MMP2, ADAMTS1, and cysteine cathepsins) indicate diminished capacity for matrix degradation, a well-established requirement for endothelial sprout invasion. ^34,36,54–60^ Together, the adhesion signature supports a model of over-stabilized adhesions coupled to inadequate local matrix remodeling, which would favor dysmorphic sprouts and failed penetration rather than cohesive, directed cords.

Consistent with impaired invasion, we observed a superimposed shift in matrix composition within the aortic secretome, with increased fibril-associated proteins (FBLN5, COL12A1),^61–63^ decreased small leucine-rich proteoglycans DCN^64^ and hyaluronan-link proteins (HAPLN1),^65^ and a transition in basement-membrane constituents (LAMA4 to LAMA5).^67,68^ Together, these changes indicate maturation toward a stabilized, less permissive matrix that disfavors sprout initiation and extension. In particular, LAMA5-enriched endothelial basement membranes characterize relatively quiescent vasculature and are less supportive of sprouting than LAMA4-rich matrices, which facilitate tip-cell programs via DLL4/Notch signaling.^66–68^ These changes suggest that sprouting is not merely inhibited but rendered structurally incompatible with the remodeled extracellular matrix and adhesion landscape.

Adhesion and invasion alterations occurred alongside cytoskeletal and trafficking changes that plausibly reinforce the migration defect observed in monolayer assays.^94,95^ A reduction in ACTR2 (Arp2/3), a core nucleator of branched actin networks that power lamellipodial protrusion,^39,40^ with concomitant reductions of the cortical spectrin scaffold SPTAN1, would further destabilize the membrane–actin cortex needed to support protrusive persistence and membrane mechanics.^41^ In smooth-muscle–enriched elements of the ring, decreased LMOD1 (an actin nucleator/organizer) and altered PPP1R12A/MYPT1 (a myosin phosphatase regulatory subunit governing actomyosin phosphorylation) indicate a shift in contractile balance that disfavors productive protrusion–retraction cycles.^96,97^ At the cortical interface, decreased AHNAK, a membrane–cytoskeleton scaffold that complexes with ANXA2/S100A10 and cortical actin, further supports loss of a robust cortex–membrane coupling needed for protrusions.^34,35^ By contrast, increased PDLIM1 (an α-actinin-binding scaffold localized to stress fibers) suggests reinforcement of stress-fiber–rich, adhesion-stabilizing architectures rather than the dynamic lamellipodial arrays required for tip-cell migration.^98^ Emerging work additionally connects PDLIM proteins to actin-dependent endosomal recycling linking this scaffold to receptor trafficking.^99^

Converging with these cytoskeletal shifts, TSST-1 seems to attenuate the endocytic–recycling axis. Decreased EHD1 would be expected to compromise β1-integrin recycling, focal-adhesion turnover, and cell spreading/migration. Mechanistic studies show EHD1 is required for integrin return from the endocytic recycling compartment.^42,43^ Decreases in the clathrin adaptor subunit AP2B1 (AP-2 β2-adaptin), a cargo/coat organizer central to clathrin-mediated endocytosis of receptors, are consistent with reduced integrin internalization and perturbed cargo selection at coated pits.^44^ Likewise, lowering COPG1 may impair Golgi–ER and intra-Golgi trafficking mediated by COPI, with potential consequences for receptor processing and recycling.^45^ Because integrin trafficking is tightly coupled to directional motility and growth-factor crosstalk, defects across these modules would be predicted to weaken directional persistence and diminish the VEGF–Notch amplification that underlies tip–stalk coordination during sprouting.

In line with the tissue-level and intracellular phenotypes outlined above, the secreted milieu of TSST-1–treated aortic rings exhibited a coordinated anti-angiogenic profile. Two potent endogenous inhibitors, THBS1 and SERPINE1, were markedly elevated. THBS1 suppress endothelial motility through CD36 engagement and by sequestering pro-angiogenic growth factors, thereby dampening VEGFR2-dependent signaling and migration.^46^ SERPINE1 constrains pericellular proteolysis by inhibiting uPA/tPA, limiting plasmin-dependent matrix degradation that is required for invasion.^47^ Concomitant increases in IGF2R, a scavenger for IGF-II, and in BMPER, a modulator capable of antagonizing BMP4 signaling in endothelium, collectively impose multilayered brakes on pro-angiogenic pathways.^48,49^ Conversely, several secreted pro-angiogenic supports were reduced, including PROS1 (linked to endothelial motility and barrier remodeling), SERPINE2/PN-1 (which, in specific contexts, preserves matrix integrity and favors angiogenesis), and LGALS3BP (reported to promote endothelial tubulogenesis).^51–53^ The concordant increase in inhibitors coupled with loss of promoters argues for active extracellular reprogramming of the angiogenic niche, rather than passive loss of angiogenic competence.

The identification of a discrete structural motif mediating TSST-1’s effects on endothelial movement represents an advancement in the conceptual understanding of SAg biology. The dodecapeptide, previously implicated in epithelial translocation and vaginal epithelial activation,^23,24^ emerged here as the critical determinant of both anti-migratory and anti-angiogenic activity. Structure–function mapping localized maximal functional impact to the coil/loop segment immediately N-terminal to the helix, as F119A abolished migration inhibition, whereas substitutions within the helical portion (S127A, T128A) only partially restored migration, indicating that helical residues contribute to but are not sufficient for full activity. In the more complex aortic ring assay, where sprouting integrates migration, proliferation, matrix remodeling, and multicellular coordination, multiple coil-region substitutions (F119A, D120A, K122A, Q123A) together with a single helical variant (I126A) restored sprouting to control levels, underscoring differential structural requirements for inhibiting two-dimensional motility versus three-dimensional angiogenesis. These data highlight the coil/loop surface as particularly critical, consistent with prior work assigning this exposed loop a role in binding non-hematopoietic cells,^23,71^ and motivate future efforts to define the relevant endothelial/perivascular binding partners.

Collectively, our findings show that TSST-1 blocks vascular repair by shutting down core drivers of endothelial migration and angiogenesis through the actions of the dodecapeptide, offering a tractable handle to disrupt SAg–endothelium interactions. This multifaceted disruption supports a model in which TSST-1 exploits endothelium plasticity, distinct from yet convergent with SEC and β-toxin, to suppress vascular repair, which could convert sites of tissue injury into persistently non-healing niches suited for *S. aureus* persistence.

### Limitations of the study

This study primarily uses *in vitro* endothelial assays and an *ex vivo* rabbit aortic ring model, which provide mechanistic insight but do not fully capture the inflammatory, hemodynamic, and multicellular complexity of *in vivo S. aureus* infection. TSST-1 concentrations were selected to enable robust mechanistic analysis and may not precisely reflect local toxin levels during infection. While proteomic and secretome profiling revealed coordinated pathway shifts, these analyses are correlative and do not establish direct causality for individual molecular changes. Finally, the endothelial receptors and signaling pathways engaged by the TSST-1 dodecapeptide remain to be defined.

## RESOURCE AVAILABILITY

### Lead contact

Requests for further information and resources should be directed to and will be fulfilled by the lead contact, Wilmara Salgado Pabón.

### Materials availability

This study did not generate new unique reagents.

### Data and cote availability

- All data reported in this paper will be shared by the lead contact upon request.
- This paper does not report original code.

## ACKNOWLEDGMENTS

We thank Darrell R. McCaslin, Ph.D. in the Biophysics Instrumentation Facility, Department of Biochemistry, University of Wisconsin−Madison for their support in CD data acquisition, analysis, and interpretation. We are grateful to Grzegorz Sabat, M.S., Proteomics Specialist, at the University of Wisconsin Biotechnology Center Mass Spectrometry Core Facility for mass spectrometry support. This work was supported by NIH grant R01AI34692-01 to W.S.-P., AHA grant 23PRE1026138 to S.S.T., NIEHS grant 1R00ES034058-05 to K.P.M.M. AVIV Model 420 Circular Dichroism Spectrometer data were obtained at the University of Wisconsin−Madison Biophysics Instrumentation Facility, which was established with support from the University of Wisconsin−Madison and grants BIR-9512577 (NSF) and S10RR013790 (NIH).

## AUTHOR CONTRIBUTIONS

Conceptualization, S.S.T and W.S.-P.; methodology, K.J.K; investigation, S.S.T, O.F.B., P.M.T., X.-J.W., A.G.C, D.S., K.P.M.M. and W.S.-P.; formal analysis, S.S.T, O.F.B., P.M.T., D.S., K.P.M.M., and W.S.-P.; visualization, S.S.T, P.M.T., A.G.C, and W.S.-P.; writing, S.S.T and W.S.-P.; project administration; W.S.-P.

## METHODS

### Media and reagents

Phenol red–free Medium 200 (M200PRF500), low serum growth supplement (LSGS; S00310), 0.025% trypsin-EDTA (R001100), VEGF (PHC9344), Pierce Protease Inhibitor Mini Tablets (A32953), Halt™ Protease Inhibitor Cocktail, EDTA-Free (78439), Dynabeads™ (10103D, 10104D), Detoxi-Gel™ (20339), penicillin-streptomycin (15140122), amphotericin B (15290018), Qubit™ Protein Broad Range kit (A50668) and human umbilical vein endothelial cells (HUVECs; C0035C; RRID: CVCL K312) were purchased from Thermofisher Scientific / Fisher Scientific.

HiTrap Heparin HP columns (17040701) were obtained from Cytiva. ToxinSensor™ LAL Endotoxin Assay Kit (L00350) was purchased from Genscript. CellTiter 96® AQueous One Solution (G3582) were purchased from Promega. Mitomycin C (M4287) was purchased from Sigma Aldrich. Proteome Profiler™ Human Angiogenesis Antibody Array (ARY007) was purchased from R&D Systems. IRDye 800CW Streptavidin (926-32230) was purchased from LI-COR. Growth factor reduced Matrigel (GFR-Matrigel; 356231) and 12 well (3513) and 48 well (3548) Costar tissue culture plates were purchased from Corning. 96 well plates (P96-1.5P) were purchased from Cellvis. Axitinib (HY-10065) was purchased from MedChemExpress. 0.1 mm glass beads (P000929LYSK0A.0) and soft tissue homogenizing kits (P000933-LYSK0-A) were purchased from Bertin Corp. 4-well culture inserts (80466) and angiogenesis slides (81506) were purchased from ibidi. Mycostrip™ Plus (LT07-701) and MycoStrip**®** (REP-MYS-10) Mycoplasma Detection Kits were purchased from Lonza and InvivoGen, respectively.

### Endothelial cell culture conditions

Immortalized human aortic endothelial cells (iHAEC) are a recently established cell line shown to retain phenotypic and functional characteristics of primary cells, serving as a large-vessel model system in which to address questions relevant to vascular biology.^17^ Cells were grown at 37°C, 5% CO_2_ in Medium 200 supplemented with LSGS, final concentrations of: FBS 2%, hydrocortisone 1 μg mL^−1^, human epidermal growth factor 10 ng mL^−1^, basic fibroblast growth factor, 3 ng mL^−1^, heparin 10 μg mL^−1^). Cells were maintained on 1% gelatin-coated plates unless otherwise stated. Cells were passaged at least twice before use in experiments. iHAECs were used at passages between 4 and 12 from a single clone. Mycoplasma-testing was conducted every 6 months using MycoAlert™ Plus (Lonza) or MycoStrip**®** (InvivoGen) Mycoplasma Detection Kit.

### Protein purification and concentrations used

TSST-1 and its respective dodecapeptide single-residue substitution mutants were expressed in *E. coli* BL21 (DE3) cells harboring a plasmid encoding N-terminal His6-tagged constructs. Cultures were initiated in 1 L LB broth (5 g yeast extract, 10 g tryptone, 10 g NaCl) containing 100 μg mL^−1^ carbenicillin and grown at 37°C to an OD600 of 0.4–0.8. Expression was induced with 1 mM IPTG and cultures maintained in the incubator shaker for 4 hours at 30°C, 250 rpm. Cultures were pelleted via centrifugation (6,000 × g, 10 min, 4°C) and resuspended in 30 mL binding buffer (20 mM NH_2_C(CH_2_OH)_3_ · HCl, 500 mM NaCl, 20 mM imidazole, pH 7.9) along with three Pierce Protease Inhibitor Mini Tablets. 10 mL aliquots were homogenized with 7 g of mm glass beads using a Precellys Cryolys Evolution homogenizer (9900 rpm, six 30 s cycles, 60 s rests, 4°C) (Bertin Corp.). Lysates were then centrifuged at 50,000 × g, 40 min, 4°C. Resulting supernatant was then manually syringe filtered through a 0.45 μm membrane. All proteins were purified by Ni²⁺-affinity chromatography on a HiTrap Heparin HP column (Cytiva) via gradient elution (20–250 mM imidazole, 50 mM NH_2_C(CH_2_OH)_3_ · HCl, 500 mM NaCl, pH 7.9) and subsequently dialyzed overnight against 4 L PBS (pH 7.4) at 4°C. Protein capture and purity were confirmed by sodium dodecyl sulfate-polyacrylamide gel electrophoresis (SDS-PAGE) followed by Coomassie blue staining to reveal a single band. Protein concentration was determined using the Qubit™ Protein Broad Range Assay Kit (Thermofisher Scientific). Endotoxin contamination was reduced using Detoxi-Gel™ endotoxin removal resin, and final endotoxin levels were measured with the ToxinSensor™ Chromogenic Limulus Amebocyte Lysate (LAL) Endotoxin Assay Kit, ensuring levels ≤0.05 ng mL^−1^. TSST-1 was used at the sub cytotoxic concentrations of 6 – 50 μg mL^−1^, depending on the sensitivity of the assay. *S. aureus* produces TSST-1 as high as μg mL^−1^ in liquid culture and 16,000 μg mL^−1^ in thin-film biofilms^100^. Septic lesions, such as vegetations, support bacterial growth as high as 5 × 10^9^ *cfu*^15^. We expect infected tissues to be exposed to these toxins at high levels.

### EdU proliferation assay

To capture dodecapeptide residue specific contributions on cell proliferation, passage 11 iHAEC cultured in Medium 200 supplemented with LSGS were seeded into a 96 well plate at 7,000 cells per well. Following 4 hours of incubation at 37°C with 5% CO₂ for cell attachment, media was gently aspirated and replaced with toxin-conditioned media (50 μg mL⁻¹), allowing for biological/technical replica per toxin condition along with media-only controls in triplicate. Cells were then incubated overnight to reach ∼80% confluency. After experimental treatments, cells were incubated with EdU (10 µM final concentration) for 2 h at 37°C in complete growth medium to label cells undergoing DNA synthesis. Cells were labeled with EdU for 10 minutes. Following EdU incorporation, cells were washed twice with phosphate-buffered saline (PBS) and fixed in 1% formaldehyde/PBS for 10 min at room temperature and washed three times in PBS. The cells were then permeabilized with 0.1% Triton X-100/PBS for 15 min. Samples were incubated for the Click reaction in a solution containing working concentrations of 2.5ug/ml 488 Azide, 2 mg/ml Sodium Ascorbate, and 2 nM Copper Sulfate in PBS. After washing, nuclear counterstaining was performed using DAPI. The plate was imaged using the ImageXpress Nano (Molecular Devices) imaging system. Images were analyzed for average integrated nuclear intensity using the MetaXpress high content image analysis software. Data were analyzed using GraphPad Prism.

### Cell-exclusion assay

Cell-exclusion assays were performed using 4-chamber silicone inserts (ibidi) to assess the effects of TSST-1 and respective dodecapeptide region mutations on the migration and proliferation of immortalized human aortic endothelial cells (iHAECs). All experiments were conducted using iHAECs between passages of 7 and 13 grown in Medium 200 supplemented with LSGS. Inserts were placed into uncoated 12-well, tissue culture-treated plates using sterile tweezers. Each chamber was subsequently seeded with 27,500 cells and incubated at 37°C with 5% CO₂ for 4 h for cell attachment. The medium was then replaced with fresh toxin-conditioned medium at 100 μg mL⁻¹, and cells were incubated overnight. Following overnight treatment, the inserts were removed. The conditioned media was collected, returned to the wells, and supplemented with additional toxin-conditioned media to achieve a final volume of 1.5 mL per well. Toxin conditions included TSST-1 (16 μg mL⁻¹), and all respective dodecapeptide residue substitutions. Mitomycin C (2 μg mL⁻¹) was also paired with each condition to isolate contributions of proliferation throughout the time course and was added immediately following insert removal. Plates were set in a Leica DMi8 microscope equipped with a Tokai Hit stage-top incubator set to 37°C and 5% CO₂ for the duration of image capture. Phase-contrast images were captured every 30 minutes over a 24-hour period using a HC PL FLUOTAR 10x/0.13 objective lens to monitor wound closure. Three independent experiments were conducted for each treatment condition, and images were analyzed automatically using ImageJ. Wound area was quantified using the Wound healing Size Tool plugin in ImageJ software.^101^ Images were processed by applying the following initial settings: Variance window radius = 50, Threshold value = 100, Percentage of saturated pixel =0.001%, with additional adjustments for individual runs to ensure accuracy of gap borders.

### Quantification of single-cell migration using CellTraxx

Single-cell migration trajectories were obtained using CellTraxx, an automated cell-tracking platform designed for high-throughput and time-resolved analysis of motile cells. Time-lapse phase–contrast images from cell-exclusion assays were acquired every 30 min for 24 h using a Leica DMi8 equipped with a Tokai Hit stage-top incubator for live cell imaging at 37°C and 5% CO₂. Raw image sequences were imported into CellTraxx, and tracking was performed using the software’s integrated centroid-detection and object-linking algorithms.^26^ Briefly, CellTraxx identifies individual cells in each frame by adaptive thresholding followed by morphological segmentation, then links these detections across frames using a predictive nearest-neighbor model to reconstruct full cell trajectories. Default tracking parameters were used. To ensure high-quality trajectories, tracks with fewer than 5 continuous frames, excessive frame gaps, or algorithm-flagged ambiguities were excluded prior to analysis. Each validated track was visually inspected within the CellTraxx interface to confirm accurate linking, especially in regions of high cell density. For each trajectory, CellTraxx automatically computed frame-to-frame displacements, instantaneous velocity, cumulative path length, Euclidean displacement from origin, and angular changes between successive steps. Instantaneous velocity (μm min⁻¹) was calculated directly from frame-to-frame displacement. Euclidean distance was defined as the straight-line distance between the starting and ending positions of each cell. Directional persistence was quantified using the forward displacement index (FDi), calculated as net displacement along the wound axis divided by cumulative path length. Confinement index, a measure of spatial restriction of movement, was computed as Euclidean distance divided by cumulative path length.

### Fluorescence Microscopy

For actin immunofluorescence, iHAECs were cultured to semi-confluence on fibronectin-coated glass slides. Cells were fixed in cold methanol, washed, permeabilized with 0.1% Tween-20, and blocked in 1% BSA. Actin filaments were stained using Acti-Stain 555 phalloidin (Cytoskeleton, Inc., Denver, CO). Slides were mounted with ProLong™ Gold Antifade containing 4’,6-diamidino-2-phenylindole (DAPI; Molecular Probes, Life Technologies, Carlsbad, CA). Fluorescence images were acquired using a Leica DMi8 upright epifluorescence microscope equipped with appropriate filter sets. Multichannel images were pseudocolored and merged using ImageJ (NIH, Bethesda, MD).

### Proteome profiler™ human angiogenesis array

iHAECs were seeded onto gelatin-coated 96-well plates at 7,000 cells per well and cultured to near confluence. Fresh media containing 50 μg mL^−1^ TSST-1 was added, and plates were incubated for 24 h at 37°C with 5% CO₂. Conditioned media was collected and stored at -80°C until analysis. Three biological replicates with three technical replicates per treatment were obtained. Relative protein expression was assessed using the Proteome Profiler™ Human Angiogenesis Antibody Array kit with fluorescent detection with procedural modifications for fluorescent analysis according to the manufacturer’s instructions. Conditioned media (120 μL) was incubated with biotinylated detection antibodies for 1 h at room temperature. During this time, capture antibody membranes were blocked with kit-provided blocking buffer. The sample-antibody mixture was then applied to membranes and incubated overnight at 4°C. Following three sequential 10 min washes, membranes were incubated with IRDye 800CW Streptavidin (1:2000) for 30 min while protected from light exposure. They were then washed again (x3) and imaged using an Azure c600 (120 μm resolution, auto-intensity). Mean pixel intensities from duplicate spots were averaged using Image Studio Software (LI-COR; RRID: SCR_013715), and fold-changes relative to untreated controls were calculated. Proteins unlikely to be produced by endothelial cells - angiopoietin-1, angiostatin/plasminogen, EG-VEGF, FGF-4, leptin, platelet factor 4, and serpin B5 – were identified through a comprehensive review and were excluded based on GTExPortal and Expression Atlas data. These analytes were not detected in iHAECs.

### *Ex-vivo* aortic ring model of sprouting angiogenesis

Juvenile mixed-sex New Zealand white rabbits (2-3 kg) were obtained from Charles River Laboratories (Massachusetts) and Western Oregon Rabbit Co. (Oregon) and housed at the Research Vivarium at the School of Veterinary Medicine, University of Wisconsin−Madison, according to established guidelines and the protocol approved by the Institutional Animal

Care and Use Committee (protocol V006634-R02). All animals were individually housed with *ad libitum* access to food and water. Rabbits were allowed at least four days for acclimatization and were clinically determined to be healthy by a veterinarian prior to euthanasia. Following euthanasia, the thoracic and abdominal aorta segments were immediately excised and placed in phosphate-buffered saline (PBS) for dissection. Excess fascia and connective tissue were removed, and 1–1.5 mm² aortic cross-sections were prepared using a scalpel. Aortic ring explants were cultured by modifying the thin-layer method.^102^ The sections were embedded in 200 μL of phenol red-free growth factor-reduced Matrigel (GFR-Matrigel, Corning) in 24-well plates. After 10 min of polymerization at 37°C in 5% CO₂, 200 μL of supplemented Medium 200 was added to each well, and the plates were incubated at 37°C in 5% CO₂ for up to 14 days.

The Medium 200 was supplemented with LSGS, 100 U mL⁻¹ penicillin-streptomycin (Thermofisher Scientific), 2.5 μg mL⁻¹ amphotericin B (Thermofisher Scientific), and relevant treatment conditions. Media (with or without treatments) was changed every 3-5 days. Images were captured after 14 days using a Leica DMi8 microscope equipped with a Tokai Hit stage-top incubator set to 37°C, 5% CO₂. A 4x/0.13 HC PL FLUOTAR objective lens was used for image acquisition, and sprouting of aortic rings was assessed for surface area in millimeters squared at day 14 using ImageJ.^103^ A minimum of three rabbits were used for each experimental condition, with at least three thoracic sections and three abdominal sections obtained for each biological replicate.

### Protein extraction and processing for mass spectrometry analysis

For protein extraction from tissue samples, aortic rings were prepared and cultured as described under the *ex vivo* explant model methodology section. Following a 14-day culture period, aorta tissues were homogenized in RIPA buffer (10 mM Tris-HCl pH 7.5, 150 mM NaCl, 0.5 mM EDTA, 0.1% SDS, 1.0% Triton X-100, 1.0% sodium deoxycholate) supplemented with Halt™ Protease Inhibitor Cocktail, EDTA-Free. For every milligram of wet tissue, 15 µL of RIPA buffer was added to 2 mL O-ring tubes containing 2.8 mm beads (1.2 g beads per 30 mg tissue).

Tissue was homogenized using a Bead Mill 4 instrument (Fisher Scientific) at 5000 rpm for 45 seconds, followed by cooling on ice for 5 min, repeated 10 times. Lysates were centrifuged at 12,000 rpm (13,680 × g) for 15 minutes at 4 °C and the supernatants collected. Protein concentrations were measured using a Qubit™ Protein Assay Kit (Invitrogen). To remove superantigens from aortic ring supernatant, magnetic bead separation was completed using Dynabeads™. 1 mg Dynabeads were washed three times with 900 µL PBS (pH 7.5), incubated in a rotator at room temperature for 2 min between washes and magnetized for 2 min at room temperature to prepare beads for sample loading. Conditioned medium from ring cultures was collected at day 14, centrifuged (12,000 × g, 15 min, 4°C), supplemented with 9 µL Halt™ Protease Inhibitor Cocktail per 900 µL, and then incubated with the prepared Dynabeads™ for 1 h at 4°C. Sample is then magnetized and supernatant removed and saved. Samples were stored at -80°C until LC-MS/MS analysis. Protein samples, including Dynabeads™ eluates, were analyzed by SDS-PAGE to confirm purity. Selected bands were excised, digested with trypsin, and desalted. Peptides were dried and prepared for LC-MS/MS analysis to identify and quantify protein abundance.

### Liquid Chromatography-Tandem Mass Spectrometry

Proteins from fresh media, tissues and supernatants from untreated controls, and TSST-1-treated conditions were submitted to the University of Wisconsin Biotechnology Center Mass Spectrometry Core Facility for Liquid Chromatography-Tandem Mass Spectrometry (LC-MS/MS). Equal amounts of proteins from three biological replicates within each treatment condition (processed separately) were pooled to generate one sample per condition. Raw data from LC-MS/MS was analyzed by UW−Madison Mass Spectrometry Facility. To ensure that only proteins originating from the aortic explants were included, culture media from both control and TSST-1 conditions were processed and analyzed in parallel, and proteins detected in fresh media alone were excluded from downstream analysis. Only proteins with total spectral counts ≥10 across all samples were retained to eliminate low-confidence identifications and stochastic peptide matches. LC-MS/MS data were searched against Oryctolagus cuniculus (rabbit) and Bos taurus (cattle) UniProt database using standard search parameters, and peptide–spectrum matches were filtered at a 1% false discovery rate using a target–decoy approach. Protein-level identifications were assembled using parsimonious inference. Proteins matching to fresh media and Matrigel components and common contaminants were eliminated. The remaining proteins in the list were *bona fide* aortic tissue proteins. Total Spectral Count Normalization (TSN) was applied to correct for sample-to-sample variation in overall mass spectrometric signal. For each sample, the spectral counts for all identified proteins were summed, and individual protein counts were divided by this total to generate normalized abundance values scaled to a common constant. These normalized values were used for downstream fold-change calculations.

### Pathway and process enrichment analysis

Pathway and process enrichment analysis was performed with the Metascape platform.^33^ For each given gene list, the Metascape platform performs pathway and process enrichment analysis with the following ontology sources: KEGG Pathway, GO Biological Processes, Reactome Gene Sets, Canonical Pathways, CORUM, WikiPathways, and PANTHER Pathway. All genes in the genome are used as the enrichment background. Terms with a *p*-value < 0.01, a minimum count of 3, and an enrichment factor > 1.5 (the enrichment factor is the ratio between the observed counts and the counts expected by chance) are collected and grouped into clusters based on their membership similarities. *p*-values are calculated based on the cumulative hypergeometric distribution and *q*-values are calculated using the Benjamini-Hochberg procedure to account for multiple testings. Kappa scores are used as the similarity metric when performing hierarchical clustering on the enriched terms, and sub-trees with a similarity of > 0.3 are considered a cluster. The most statistically significant term within a cluster was chosen to represent the cluster. The Metascape platform identified the top 100 enriched pathways and processes. A detailed list of the top 20 pathways and processes and their protein assignments are listed in Table S1 and S2.

### TSST-1 dodecapeptide expression vector construction

All constructs were generated in the pET25bHSVdelTEV plasmid backbone and cloned into *Escherichia coli* DH5α before being transformed into *E. coli* BL21 (DE3). Unless otherwise noted, reagents were used as supplied by the manufacturer and incubations were performed at recommended temperatures and times. *tstH* was PCR-amplified from *S. aureus* strain MN8 with Phusion high-fidelity DNA polymerase (New England Biolabs) using primers tsthNdeIF and tsthpET25bXhoIR. The *tstH* PCR fragment and pET25bHSVdelTEV backbone were digested with NdeI and XhoI and ligated with T4 DNA ligase (New England Biolabs) creating the plasmid pET25bHSVdelTEV_tstH. Site-directed mutants were generated in the pET25bHSVdelTEV_tstH backbone with the QuikChange II Site-Directed Mutagenesis Kit (Agilent Technologies) following the manufacturer’s protocol. All coding sequences and vector–insert junctions were confirmed by Sanger sequencing using the T7_prom primer. All primers were purchased from Integrated DNA Technologies (Supplementary Table 1).

### Circular Dichroism Spectroscopy

All proteins were expressed and purified to homogeneity and submitted to the University of Wisconsin Biophysics Instrumentation Facility. Because each mutation replaces only a single residue and does not alter aromatic content, the molecular mass used for all variants was set to 26,473 Da (234 total amino acids). Aromatic composition (3 Trp and 9 Tyr) is unchanged across mutants; thus, molar extinction coefficients were calculated uniformly using the Pace et al. parameters, yielding ε280 = 29,910 M⁻¹ cm⁻¹ with no contributions from disulfide bonds. Protein concentrations were quantified by absorbance at 280 nm. When low-intensity scattering tails were observed in the UV spectrum (despite prior centrifugation) no further light-scattering correction was applied, consistent with our use of measured A280 values for all concentration calculations. UV absorbance spectra were recorded at 25°C using self-masking quartz cuvettes of 0.4 cm width and 1.0 cm pathlength on a Shimadzu UV2700 instrument (0.2 nm bandpass).

Immediately after UV measurement, the same samples were transferred to 0.1 cm pathlength quartz cuvettes for CD spectroscopy. CD spectra were acquired at 25°C on an Aviv 420 spectropolarimeter using a 1.0 nm bandpass. Spectra from PBS buffer were collected under identical conditions and subtracted from all samples. Because of buffer absorbance, usable CD wavelengths were limited to ≥200 nm. Ellipticity values were converted to molar ellipticity using concentrations in µM and a residue count of 233 residues (one fewer than the full protein length).

### Statistical Analysis

Statistical analyses were conducted using GraphPad Prism software (RRID: SCR_002798). For each experiment, the precision measures, as well as the number of technical and biological replicates, are detailed in the figure legends. Information on the number of cells analyzed, measurements taken, and experiment timing can be found in the Method Details section for each experimental setup. Aortic ring sprouting data were analyzed using a one-way ANOVA with Holm-Šídák’s multiple comparisons test, with a significance level of α = 0.05. Cell-exclusion assay data were analyzed using two-way ANOVA with Holm-Šídák’s multiple comparisons test, while EdU cell proliferation was evaluated via non-parametric ANOVA. Statistical significance with α = 0.05 is indicated as follows: * *p* < 0.05, ** *p* < 0.01, *** *p* < 0.001, and **** *p* < 0.0001 (GraphPad Prism 11.0.0).

