## Supplementary material for "A TSST-1 structural motif disrupts endothelial programs required for vascular regeneration": Table S5

Supplementary Table 5

| Name | Sequence (5'–3') |
|---|---|
| tsthNdeF | ttacCATATGTCTACAAACGATAATATAAAGG |
| tsthpET25bXhoIR | ttacCTCGAGATTAATTTCTGCTTCTATAGTTT |
| T7_prom | taatacgactcactataggg |
| tsthF119Afor | ccaaagGCAgataaaaaacaattagctatatcaacttta |
| tsthF119Arev | ttgtttttatcTGCctttggccaatactttaa |
| tsthD120Afor | aagttcGCAaaaaaacaattagctatatcaacttta |
| tsthD120Arev | taattgtttttTGCgaactttggccaatactt |
| tsthK121Afor | ttcgatGCAaaacaattagctatatcaactttagacttt |
| tsthK121Arev | agctaattgtttTGCatcgaactttggccaata |
| tsthK122Afor | gataaaGCAcaattagctatatcaactttagactttgaa |
| tsthK122Arev | tatagctaattgTGCtttatcgaactttggcca |
| tsthQ123Afor | taaaaaaGCAttagctatatcaactttagactttgaaatt |
| tsthQ123Arev | tgatatagctaaTGCtttttatcgaactttgg |
| tsthL124Afor | aaacaaGCAgctatatcaactttagactttgaaattcgt |
| tsthL124Arev | agttgatatagcTGCttgtttttatcgaactt |
| tsthA125Sfor | caattaTCAatatcaactttagactttgaaattcgtcat |
| tsthA125Srev | taaagttgatatTGAtaattgtttttatcgaa |
| tsthI126Afor | ttagctGCAtcaactttagactttgaaattcgtcatcag |
| tsthI126Arev | gtctaaagttgaTGCagctaattgtttttatc |
| tsthS127Afor | gctataGCAactttagactttgaaattcgtcatcagcta |
| tsthS127Arev | aaagtctaaagtTGCtatagctaattgtttttta |
| tsthT128Afor | atatcaGCAttagactttgaaattcgtcatcagctaact |
| tsthT128Arev | ttcaaagtctaaTGCtgatatagctaattgttt |
| tsthL129Afor | tcaactGCAgactttgaaattcgtcatcagctaactca |
| tsthL129Arev | aatttcaaagtcTGCagttgatatagctaattg |
| tsthD130Afor | actttaGCAtttgaaattcgtcatcagctaactcaaata |
| tsthD130Arev | acgaatttcaaaTGCtaaagttgatatagctaa |
